# Relation is an option for processing context information

**DOI:** 10.1101/2022.04.14.488336

**Authors:** Kazunori D Yamada, M. Samy Baladram, Fangzhou Lin

**Affiliations:** Graduate School of Information Sciences, Tohoku University, Sendai, Japan; Artificial Intelligence Research Center, National Institute of Advanced Industrial Science and Technology, Tokyo, Japan

**Keywords:** Transformer, attention, artificial intelligence, time complexity, multilayer perceptron

## Abstract

Attention mechanisms are one of the most frequently used architectures in the development of artificial intelligence because they can process contextual information efficiently. Various artificial intelligence mechanisms such as transformer for processing natural language, image data, etc. include the attention mechanism. Since attention is a powerful component to realize artificial intelligence, various improvements have been made to enhance its performance. The time complexity of attention depends on the square of input sequence length. Methods to improve time complexity of attention are one of the hottest topics in various studies. In this study, we have devised a mechanism called relation that can understand the context information of sequential data without using attention. Relation is very simple to implement and its time complexity depends only on the length of the sequences; a comparison of the performance of neural networks with relation and attention mechanisms on several benchmark datasets shows that relation achieved the same context processing capability as attention with less computation time. Hence, relation is an ideal option for processing context information.

## 1 Introduction

Attention is a mechanism developed in 2017 to reveal the relationship between different positions in sequential data [1]. The basic principles of attention have been implemented in a variety of research fields and it exhibits outstanding performance to date. Its performance has been particularly successful in the field of natural language processing where many pre-trained models such as Bidirectional Encoder Representations from Transformers are built on the basis of attention.

Before the advent of attention, recurrent neural networks (RNNs), such as long short-term memory and gated recurrent unit were mainly used to process sequence data [2, 3]. One of the advantages of attention over RNNs is its computational speed. RNNs, due to their structure, process the tokens in the sequence in a sequential manner, which is very disadvantageous in terms of parallelization of computation. Therefore, the abundant computing resources could not be utilized efficiently. However, sequence processing using attention does not include step-by-step processing, thus allowing complete utilization of the performance computer. Since the size of data handled by deep learning methods is very large, the efficiency of computation is an important indicator for architecture selection.

This study discusses only self-attention, which is used to calculate the relationship between tokens in a single sequence. Additionally, time complexity of attention has been discussed. It is defined as *O*(*N*^2^) when the length of the input sequence is *N*. However, the data in the real world is sometimes so long that the computation time required to calculate the square of *N* can be troublesome. Therefore, many attempts have been made from various perspectives to reduce the amount of time complexity [4]. Among the methods developed to date, linear attention provides the best time complexity in which the value is reduced to *O*(*N*) [5].

Attention, including linear attention, is a very powerful method for processing contextual information in sequential data. However, according to the universal approximation theorem, any multilayer perceptron (MLP) with sufficient expressive power can approximate any nonlinear function in real world even without complex architectures, such as RNNs, convolutional networks, and attention networks [6]. In other words, attention mechanism is not necessarily an essential structure for artificial intelligence to understand the contextual information in sequences. This is indicated in one of the previous studies as well. [7]. During contextual processing using attention, the MLP is calculated for each token. This structure is called pointwise MLP. This pointwise computation is done for each token; however, it does not convey the relationship between tokens in sequences to the downstream network. Hence, attention is used for adding information between tokens in input sequences to the pointwise MLP. For a neural network to efficiently process and understand context, it needs to efficiently convey the contextual information of the entire input sequence to the downstream network. Conversely, if each pointwise MLP can appropriately add contextual information derived from the entire input sequence, it is possible to employ alternative methods without using attention.

In this study, we have devised an alternative method known as relation for conveying contextual information to each pointwise MLP. It is a simple structure that conveys contextual information from an input sequence to each pointwise MLP. The time complexity of relation is *O*(*N*), which is the same as that of linear attention. Additionally, we have analyzed if relation can replace attention and linear attention using several well-known natural language processing benchmark datasets.

## 2 Method

### 2.1 Attention and Relation

#### 2.1.1 Attention and linear attention

In this study, we have demonstrated the computation of attention *A* and linear attention *L* for the input sequence *x* ∈ ℝ^*N×m*^. The length of the input sequence is *N*. In addition, let *m* be the size of the feature vector of each token in the input sequence. First, the query, key, and value matrices are computed using the following equations:

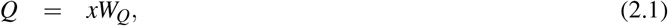

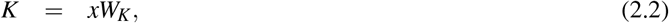

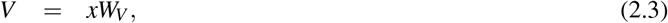

where *W*_*Q*_ ∈ ℝ^*m×d*^, *W*_*K*_ ∈ ℝ^*m×d*^, and *W*_*V*_ ∈ ℝ^*m×d*^ are the *m × d* of the trainable parameter matrix that is responsible for projecting each token of the input sequence into a vector of *d* elements where *d* is the size of attention and is called depth. Attention *A* is calculated using the following equation using the softmax function ϕ:

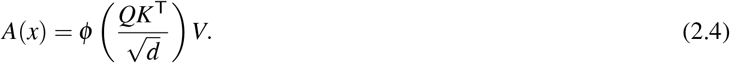

Next, linear attention *L* is computed by the following equation:

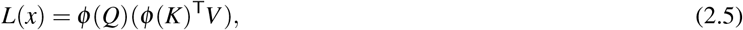

where ϕ is not a softmax function; however, it is defined by the following equation:

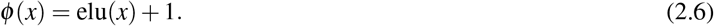

#### 2.1.2 Relation

In this study, we have defined a method known as relation for conveying the whole contextual information of the input sequence to the downstream pointwise MLP without the use of attention. First, *G* and *H* are generated using the following equations:

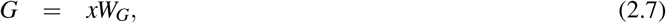

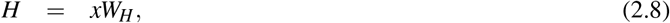

where *W*_*G*_ ∈ ℝ^*m×d*^ and *W*_*H*_ ∈ ℝ^*m×d*^ are trainable parameter matrices in *m × d* where each token in the input sequence is projected to a vector of *d* elements. Each row of *H* is considered a vector as follows:

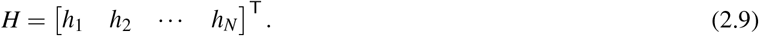

*h*^*′*^ is computed from the elementwise mean of *h*_*i*_ using the following formula:

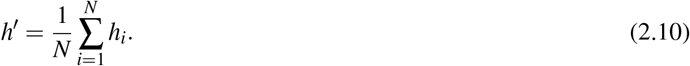

Let *H*^*′*^ be the matrix of *N* rows each row of which is the vector *h*^*′*^:

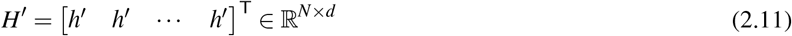

Finally, the relation *R* is calculated using the trainable parameter matrix *W ∈* ℝ^*d×d*^ as follows:

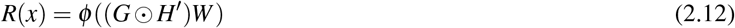

Where ⊙ is the operator for computing the Hadamard product and *ϕ* is a nonlinear activation function such as rectified linear unit.

In this relation system, *h*^*′*^ is the data containing the features of the entire input sequence. We have interpreted each row of *G* as a weight that allows each token in the input sequence to extract information from *h*^*′*^. The overall view of the relation system is shown in Figure 1.

**Fig. 1:**
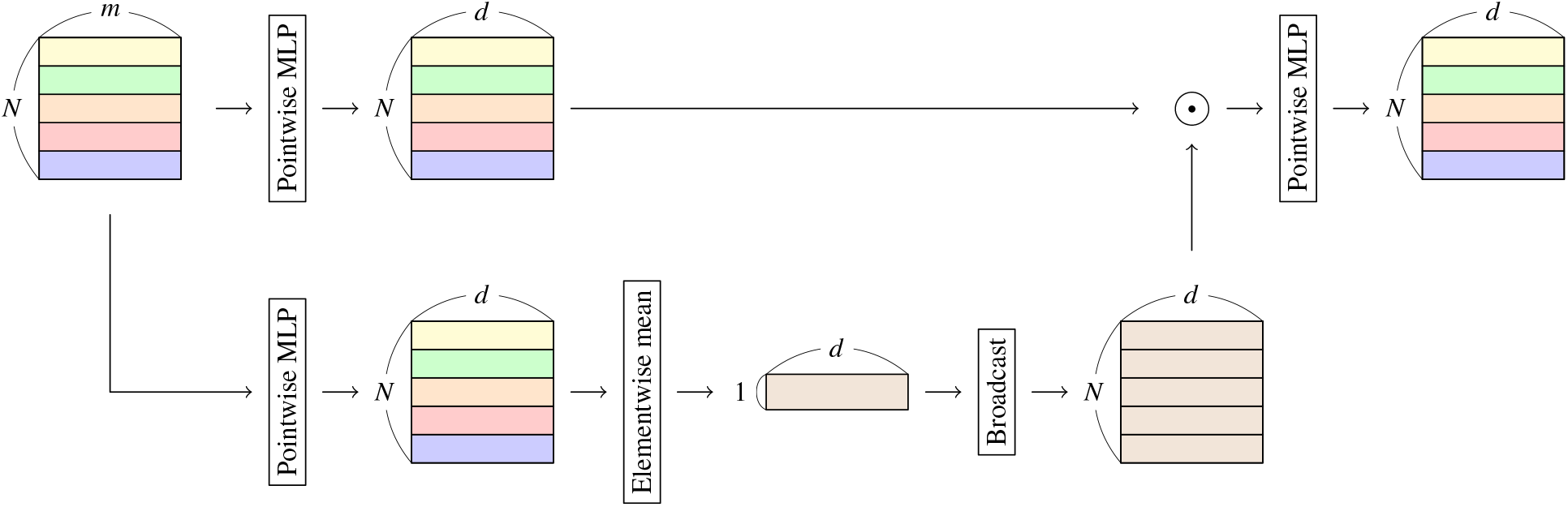
Structure of relation system. Input data is a matrix *∈* ℝ^*N×m*^ where length and feature vector size of input data is *N* and *m*, respectively. Output data is a matrix *∈* ℝ^*N×d*^ where *d* is the depth of the relation system.

### 2.2 Benchmark

#### 2.2.1 Network architecture

For all the benchmarks defined in the study, we have used a simple structure without piling up the layers to compare networks with attention, relation, and without attention or relation. In all the network structures, either attention or relation was computed from the input sequence followed by a single layer of pointwise MLP. The feature vector corresponding to the first token in the output from the pointwise MLP was used to compute the final output; the feature vector was used as input to a softmax or sigmoid function depending on the given problem (classification or regression) to produce the final result. In this study, a simple pointwise MLP without attention or relation was used as a baseline model (Baseline) for comparison. Three layers of pointwise MLP were used to ensure that the parameter size is consistent with the other networks. Details of all the networks are shown in Section *Data Availability Statement* on GitHub.

#### 2.2.2 Performance to process context

We have used General Language Understanding Evaluation (GLUE) benchmark dataset to evaluate the ability of each neural network to process contextual information. GLUE consists of a total of 11 test datasets of various types related to natural language processing (CoLA, SST-2, MRPC, STS-B, QQP, MNLI-m, MNLI-mm, QNLI, RTE, WNLI, and AX). It is the most well-known benchmark system for evaluating artificial intelligence dealing with contextual information [8]. Using the provided training dataset, the predictors were grown and the final predictor was obtained by early stopping with the patience set to 5. The constructed predictor solved the test problem and results were sent to the GLUE server to obtain GLUE scores. Scores for each test are as given in the cited paper and are not explained here. Positional encoding similar to that of a transformer was implemented [1]. Token embedding was done with a vector of 100 lengths, and global vector (GloVe) was used as the initializer for embedding [9].

#### 2.2.3 Computation time

The longest sequence of data in the GLUE dataset is 467, while that in the IMDb dataset and Reuters newswire classification dataset (Reuters) provided by TensorFlow is 2494 and 2376, respectively, which are very long [10]. As mentioned above, the time complexity of attention is *O*(*N*^2^) and that of Relation is *O*(*N*). We benchmarked the computation times on these datasets to determine the difference in actual computation time when using long input sequences. We measured the time taken for the accuracy of the validation dataset to reach 0.8 consecutively 10 times.

## 3. Results

### 3.1 Performance to Process Context

The results of GLUE-based prediction performance benchmarks are shown in Table 1. It can be observed that the prediction performance is not as good as that of existing state-of-the-art methods or complex structures such as transformer. However, the objective of this experiment is to verify if relation can process contextual information similar to attention and not whether relation can achieve high prediction performance when used for complex structures such as transformer. The GLUE score measures accuracy, correlation coefficient, or F1 score, and a high value indicates good performance. The GLUE benchmark consists of 11 evaluation items, the most important of which is AX. For the remaining 10, training and validation datasets are provided to train the predictors to make predictions on the test dataset; however, for AX, neither the training dataset nor the validation dataset are provided. AX is a problem that considers two sentences as input and classifies them into three classes. Since MNLI is a similar problem, we have used a predictor trained on this dataset to predict AX, which is completely independent of the other 10 datasets.

**Table 1:** Benchmark results with GLUE. The number in parentheses stands for the depth of attention, relation, or MLP. The best value in each block is highlighted in bold face.

| Method | All | CoLA | SST-2 | MRPC | STS-B | QQP | MNLI-m | MNLI-mm | QNLI | RTE | WNLI | AX |
| --- | --- | --- | --- | --- | --- | --- | --- | --- | --- | --- | --- | --- |
| Baseline (64) | 43.7 | 0.0 | 61.1 | **79.9/66.5** | 12.9/16.1 | 13.0/**81.0** | 38.0 | 38.0 | 52.6 | 49.3 | 57.5 | 0.7 |
| Attention (64) | 51.2 | 0.0 | 73.1 | 78.2/67.7 | 25.9/26.5 | 52.3/78.6 | 51.5 | 51.6 | 56.9 | 50.6 | 63.7 | 5.2 |
| Linear Attention (64) | 53.4 | 8.4 | 80.3 | 78.7/**69.5** | 26.7/27.4 | 49.3/76.1 | 54.5 | 52.8 | 57.2 | 52.0 | **65.1** | 5.5 |
| Relation (64) | **53.8** | **10.6** | **80.5** | 76.1/67.3 | **28.7/27.6** | 50.6/75.7 | **55.4** | **53.9** | **57.8** | **52.3** | **65.1** | **9.7** |
| Baseline (128) | 43.5 | -1.2 | 61.2 | 76.9/64.0 | 14.6/15.4 | 31.1/76.6 | 38.0 | 39.1 | 52.5 | 50.5 | 50.7 | 2.9 |
| Attention (128) | 52.0 | **10.9** | 73.5 | 77.0/67.0 | 25.0/24.0 | 50.9/**79.5** | 53.8 | 52.3 | 56.6 | 52.0 | 60.3 | 4.4 |
| Linear Attention (128) | 54.0 | 9.6 | 80.0 | **77.9/68.6** | 31.5/30.6 | 49.4/75.6 | 56.3 | 54.7 | 56.7 | **52.7** | 64.4 | 8.6 |
| Relation (128) | **54.2** | 8.7 | **80.5** | 75.1/66.3 | **32.2/31.7** | **51.6/77.6** | **56.4** | **55.1** | **58.3** | 52.1 | **65.1** | **10.2** |
| Baseline (256) | 43.8 | -1.2 | 61.7 | 76.4/63.5 | 11.9/13.8 | 3.0/82.2 | 37.7 | 38.3 | 53.0 | 52.0 | **65.1** | 0.7 |
| Attention (256) | 52.1 | 6.4 | 74.0 | 77.0/67.1 | 25.2/23.8 | 51.4/**79.2** | 53.8 | 52.3 | 56.7 | 51.9 | **65.1** | 5.1 |
| Linear Attention (256) | 54.2 | 10.7 | 80.6 | 76.4/67.8 | 31.5/30.0 | 49.9/76.0 | 57.1 | 55.3 | 56.7 | **53.3** | **65.1** | 10.0 |
| Relation (256) | **55.3** | **11.3** | **81.6** | **78.0/68.5** | **33.2/31.5** | **52.1/78.2** | **57.9** | **56.8** | **58.8** | 53.1 | **65.1** | **14.7** |

It was observed that baseline performed worse than attention or relation predictors on all datasets at all depths and very poorly on AX. Baseline is a method that does not consider any context of the input sequences. Since the parameter size of each method at each depth is unified, there is no advantage or disadvantage depending on the size of the network; the GLUE benchmark performs better when the context is taken into account; hence, the inability of baseline to consider context may be the reason for its low prediction performance. The performance of networks with attention and linear attention including the value of AX improved with increasing depth. In addition, the prediction performance using these networks was clearly better than that of baseline because the context of the input sequences is taken into account. In contrast, the network with relation also exhibited improved prediction performance depending on the size of the depth and was comparable to attention and linear attention on all datasets including AX. Thus, it is clear that contextual information can be processed by relation in the same manner as attention and linear attention.

### 3.2 Computation Time

The results of evaluating the computation time of each method using IMDb and Reuters dataset are shown in Table 2. The calculation with baseline, which consists of only pointwise MLPs could not be completed because, its accuracy did not increase with an increase in epochs. The accuracy using baseline is approximately 0.5 on IMDb and 0.3 on Reuters from the first epoch to the end of the computation. Baseline exhibited decent prediction performance on some of the GLUE datasets. Because GLUE has many short input sequences, if specific tokens are used for prediction, then is likely that in some cases good predictions can be made without considering the context. However, the sentences in IMDb and Reuters are long and composed of a variety of words; hence, it is assumed that the correct answer could not be derived without considering the context.

**Table 2:** Computation time (s) of learning phase for IMDb and Reuters. Total learning time and time per epoch is presented. The number in parentheses stands for the depth of attention, relation, or MLP. The best value in each block is highlighted in bold face.

| Method | IMDb |  | Reuters |  |
| --- | --- | --- | --- | --- |
|  | Total time | Time / epoch | Total time | Time / epoch |
| Baseline (64) | n/a | n/a | n/a | n/a |
| Attention (64) | 337 | 15.3 | 983 | 4.37 |
| Linear Attention (64) | 70.6 | 2.14 | 228 | 0.721 |
| Relation (64) | **33.0** | **1.74** | **136** | **0.591** |
| Baseline (128) | n/a | n/a | n/a | n/a |
| Attention (128) | 344 | 17.2 | 558 | 4.94 |
| Linear Attention (128) | 111 | 3.83 | 174 | 1.27 |
| Relation (128) | **54.7** | **2.88** | **111** | **0.998** |
| Baseline (256) | n/a | n/a | n/a | n/a |
| Attention (256) | 147 | 6.14 | 440 | 5.71 |
| Linear Attention (256) | 424 | 22.3 | 199 | 2.11 |
| Relation (256) | **86.1** | **4.53** | **139** | **1.61** |

The network with relation exhibited the best performance in terms of total computation time and computation time per epoch at all depths. The overall parameter size for each network is unified for each depth. As mentioned earlier, the time complexity for both linear attention and relation against the sequence length is *O*(*N*). However, when projection and depth of the input sequence are considered for calculation, the time complexity for linear attention is 2*Nd*^2^ + 3*Ndm* and relation is *Nd*^2^ + 2*Ndm* + *Nd*. Therefore, linear attention calculation requires *Nd*^2^ + *Nd*(*m −* 1) more computations. The difference in time is due to the difference in the experimental results. Thus, relation can reach the same level of prediction performance as attention and linear attention in a shorter duration.

## 4. Discussion

In this study, we devised a mechanism that considers attention and context information into account and compared its performance to networks with attention and simple MLPs without context processing capability. Attention is an excellent method used as a component of various networks including transformer. The motivation behind developing the proposed method was to improve the computation time for attention. This improvement affects various fields, such as natural language processing and image processing. Therefore, extensive research has been conducted on the improvement of computation time for attention [4]. The results indicated that time complexity of attention was *O*(*N*^2^) and that of linear attention was *O*(*N*). Thus, linear attention exhibits better performance, and our goal in this study was to create a mechanism with the same level of computational complexity and prediction performance as linear attention.

Over the years, the amount of data available has increased. The real world contains a wide variety of data, including natural language, images, biological strings, and economic time series data as well as contextual information. Relation is a method for dealing with such data. In this study, we implemented a simple approach in which the information of each token in the input sequence was aggregated into a single vector by computing an elementwise mean value of each vector projected from each token in the input sequence. Another approach was to process contextual information by aggregating it into a vector and then returning the information to the network corresponding to each token; That is, MLP alone was used instead of computations. However, the idea of MLP is not novel and its application has already been clarified by the universal approximation theorem. One of its implementations is MLP-Mixer [7]. Since the goal of this study was to improve time complexity of attention, we did not consider relation for comparison, which has a time complexity of *O*(*N*^2^), because there are better ways of capturing the entire contextual information than relation or the proposed method. In the future, we intend to continue our efforts to find new methods.

Benchmarks with GLUE, a benchmark system for natural language processing, and benchmarks on datasets containing long sequences revealed that relation can process contextual information in improved computation time. The objective of this study was to verify if relation could process context information faster than attention and not if it could achieve high prediction performance when used with complex structures such as transformer. In other words, it is unclear whether Relation will perform efficiently when used as a component of a larger network structure such as transformer. This is a limitation of this study. This, this study revealed that relation can process contextual information that a simple MLP (Baseline) could not process; however, in a small network structure it can process as much as attention. In the future, we will analyze a large network using our current computational resources. However, since relation can process context information, at least in small structures, and is computationally superior to attention, it may be a viable alternative to attention in situations where larger data must be processed within a limited time frame.

## Acknowledgement

All text in the article was translated from Japanese to English using DeepL.

## Availability

The code for this study is available in GitHub (github.com:yamada-kd/Relation.git).

